# Conjugative delivery of a miniature Cas nuclease for broad range recipient gene editing

**DOI:** 10.64898/2026.11.12.766323

**Authors:** Josephine Hotten, Dolores L. Guzmán-Herrador, Matxalen Llosa

## Abstract

Efficient delivery of the Cas endonuclease into target cells avoiding toxic effects remains one of the limiting steps of CRISPR-Cas gene editing technologies. Bacterial conjugation delivers protein and DNA directly into recipient bacteria, since the conjugative relaxase travels through a type IV secretion system covalently attached to the transferred DNA strand. We previously showed that fusion proteins relaxase- Cas12a or relaxase- base editor are translocated by conjugation and recover gRNA-guided nuclease or base-editing activity in recipient bacteria, despite the large size of the fusion. Aiming to improve the efficiency and to broaden the range of susceptible recipient bacteria, here we test translocation of the miniature type V-U4 nuclease Cas12n, fused to two unrelated conjugative relaxases, MobA and TrwC from RSF1010 and R388 respectively. Both fusions of 139 and 168 kDa were stably produced in *Escherichia coli* donor cells, mobilized their cognate plasmid as efficiently as the wild-type relaxases, and directed guide-dependent cleavage of a chromosomal target in the donor. Translocation and activity in recipient cells were tested with a lethality assay based on the loss of transconjugants viability caused by double-strand breaks introduced by the translocated nuclease. When MobA-Cas12n was delivered together with a mobilizable plasmid which encoded the guide RNA, conjugation frequency was reduced by more than one order of magnitude relative to both a non-targeting guide RNA and a nuclease-free relaxase control, indicating target-specific cleavage of the recipient chromosome. When assayed in parallel with previously characterized TrwC-Cas12a, MobA-Cas12n showed better protein yields and higher activity in recipient cells. This work extends the range of Cas proteins deliverable by conjugative relaxases to miniature nucleases. The fusion with relaxase MobA allows the use of the promiscuous RP4 conjugative system for conjugative delivery, extending the range of recipient bacteria potentially targeted by this tool.

## 1. Introduction

Bacteria are used for many biotechnological applications, from bioremediation and wastewater treatment to the production of pharmaceuticals, secondary metabolites and biofuels, or the on-site delivery of pharmaceutical compounds. Most of these applications require genetic modification of the producer strain [1]. Among the available genetic tools, CRISPR-Cas nucleases and their engineered derivatives have made target selection straightforward. These systems rely on the site-specific activity of an effector Cas protein guided to its target by a guide RNA (gRNA), and depending on the derivative used the outcome ranges from specific cleavage to base or prime editing [2].

The ability to engineer bacteria depends on the delivery of the editing tools to the target cell [3]. Most wild-type isolates, which often possess the greatest biotechnological potential, are frequently recalcitrant to transformation. Bacterial conjugation is a valuable alternative to reach a broad range of recipient bacteria. Sending the c*as* gene, however, requires fine tuning of *cas* expression in the target cell, since overexpression of *cas* genes often produces toxicity and off-target activity [4]. This tuning is especially challenging in non-model hosts, which frequently lack characterized expression systems and replicative plasmids [5]. Alternative methods, such as the delivery of a preassembled ribonucleoprotein, would bypass the expression bottleneck, but in bacteria this technique remains largely unexplored.

As an alternative, our group has developed a new strategy (hereafter referred to as ConCasTool) to deliver the desired Cas proteins, such as Cas12a or base editors, by fusing them to a conjugative relaxase. During bacterial conjugation, the conjugative relaxase cleaves the origin of transfer (*oriT*) of the transferable plasmid, remains covalently bound to the 5′ end of the transferred strand, and is translocated by a Type IV secretion system (T4SS) into the recipient cell, where it re-circularizes the DNA [6]. Thus, the Cas protein fused to the relaxase is also translocated through the conjugative T4SS, which recruits and translocates the conjugative relaxase moiety [7]. Fig 1 outlines the ConCasTool design. The donor cell carries three elements: a conjugative plasmid which provides the T4SS but lacks a functional relaxase gene; a non-mobilizable plasmid expressing the relaxase-Cas12a fusion; and a mobilizable plasmid carrying the corresponding *oriT* and encoding the gRNA. With this design, no prior genetic manipulation of the target cell is required, and Cas is delivered as a protein, so no *cas* gene has to be expressed in it. In our previous work, the nuclease Cas12a was fused to the relaxase TrwC of plasmid R388. The fusion protein was translocated through the R388 T4SS and recovered guide-dependent nuclease activity in recipient *E. coli* after translocation. ConCasTool was then extended to base editors, obtaining site-specific edition of chromosomal and plasmid genes in the recipient (Guzman-Herrador et al, 2024). In spite of its potential, a bottleneck of ConCasTool is its low efficiency, which jeopardizes its future application to edit wild type bacteria, for which conjugation efficiencies are usually low.

**Fig 1.**
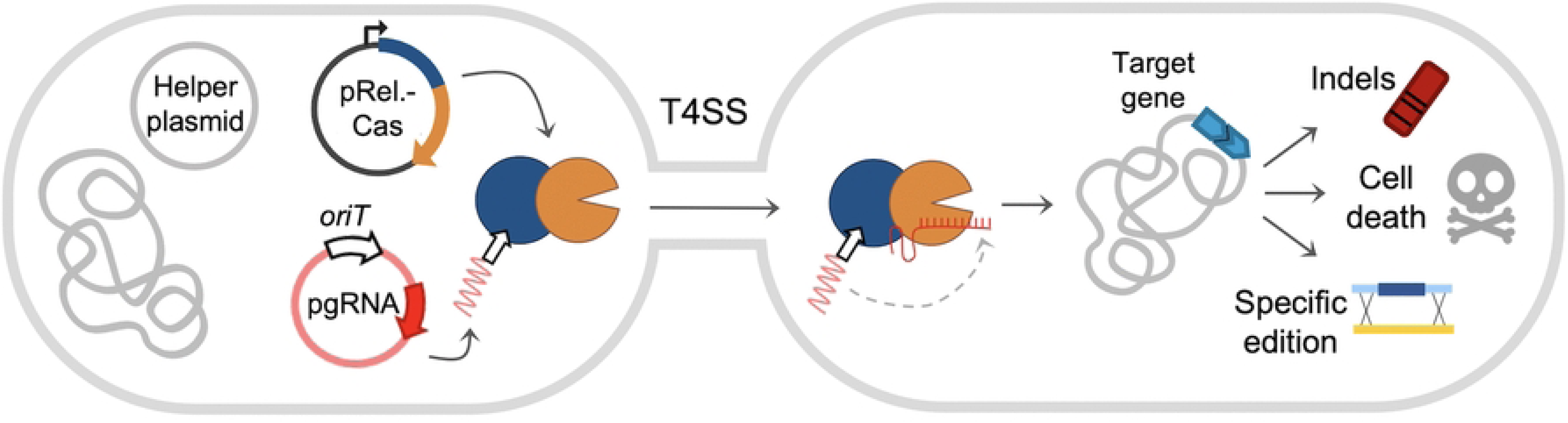
The ConCasTool rationale. The donor cell carries a non-mobilizable plasmid encoding the relaxase-nuclease fusion, a conjugative helper plasmid, and a mobilizable plasmid encoding the gRNA. The relaxase domain of the fusion protein nicks the mobilizable plasmid at the *oriT*, and the resulting nucleoprotein complex is translocated through the T4SS into the recipient cell, where the fusion protein refolds and regains Cas activity which, guided by the sgRNA to its chromosomal target, induces the double-stranded break, producing the desired editing effect.

Two of the likely limitations for the efficiency of ConCasTool are the amount of fusion protein production, and the efficient refolding in the recipient, since the relaxase has to be unfolded and refolded during this translocation process [8,9]. Relaxase-Cas fusion proteins are more than 200 kDa in size, and we observed extensive protein degradation in the donor, so that only a fraction of the protein made is available for translocation; we reasoned that smaller proteins could partially alleviate this problem. As for the refolding, it is impossible to anticipate how proteins not designed to be translocated through a T4SS are going to behave; however, it has been suggested that primitive proteins, which rely less on host chaperones, have higher capacity to refold [10].

Since CRISPR-Cas system were described, many different versions of Cas protein have been discovered or engineered. Of special interest here are the miniature members of the Cas12 family such as Cas12n, a type V-U4 effector of 506 residues that probably represents the earliest evolutionary intermediate between the transposon-associated protein TnpB and the larger Cas12 proteins [11]. This protein carries a single RuvC domain and recognizes a 5′-AAN PAM and generates staggered cuts. Double strand breaks cannot be repaired in most bacteria [12], so the targeted nuclease activity of Cas proteins usually leads to cell death. The AcCas12n from *Actinomadura craniellae* was tested in *Klebsiella pneumoniae* for specific killing, obtaining efficiencies close to 100%.

In this work, we have expanded the previous ConCasTools by developing ConCasMin, a version based on miniature Cas nucleases, with the aim of increasing the number of Cas proteins available for conjugative delivery and for addressing the low targeting efficiency that limited the original ConCas system. In addition, we aimed to extend the range of potential recipient cells which could be edited by the ConCasTools; this range is defined by the ability of the relaxase to be recruited by different T4SS. Thus, we have fused the miniature endonuclease Cas12n to the conjugative relaxases TrwC and MobA. TrwC is the relaxase of the conjugative plasmid R388, the system we previously used to deliver Cas12a. MobA is the relaxase of the mobilizable plasmid RSF1010, a highly promiscuous system that is mobilized by the T4SS of many unrelated conjugative plasmids; in particular, it can use the T4SS of plasmid RP4, whose transfer range extends beyond the Gram-negative bacteria to include Gram-positive genera of biotechnological interest [13–15]. Both fusions were produced under inducible conditions and retained the activity of their two moieties, relaxase and site-specific nuclease. To detect their translocation and activity in recipients, MobA-Cas12n endonuclease activity was detected by the drop in transconjugant viability. Thus, ConCasMin represents a promising new ConCas system which can be delivered by conjugation to a potentially broad range of bacterial recipients, including Gram-positive bacteria.

## 2. Material and methods

### 2.1. Bacterial strains

The *Escherichia coli* strains used in this work are listed in Table 1. *E. coli* strain D1210 was used for standard cloning. The particular *E. coli* strains used as donors and recipients in conjugation complementation and protein translocation, are described for each assay. The strains were grown at 37°C in Luria Bertani (LB) broth, supplemented with agar for solid culture. Antibiotics and gene expression inducers were used at the following concentrations: ampicillin (Ap) 100 µg/mL, kanamycin monosulfate (Km) 20 µg/ml, streptomycin (Sm) 300 µg/ml, gentamycin sulfate (Gm) 10 µg/mL, nalidixic acid (Nx) 20 µg/mL, anhydrotetracycline hydrochloride (aTc) 200 ng/mL, Isopropyl beta-d-1-thiogalactopyranoside (IPTG) 500 µM.

**Table 1:**
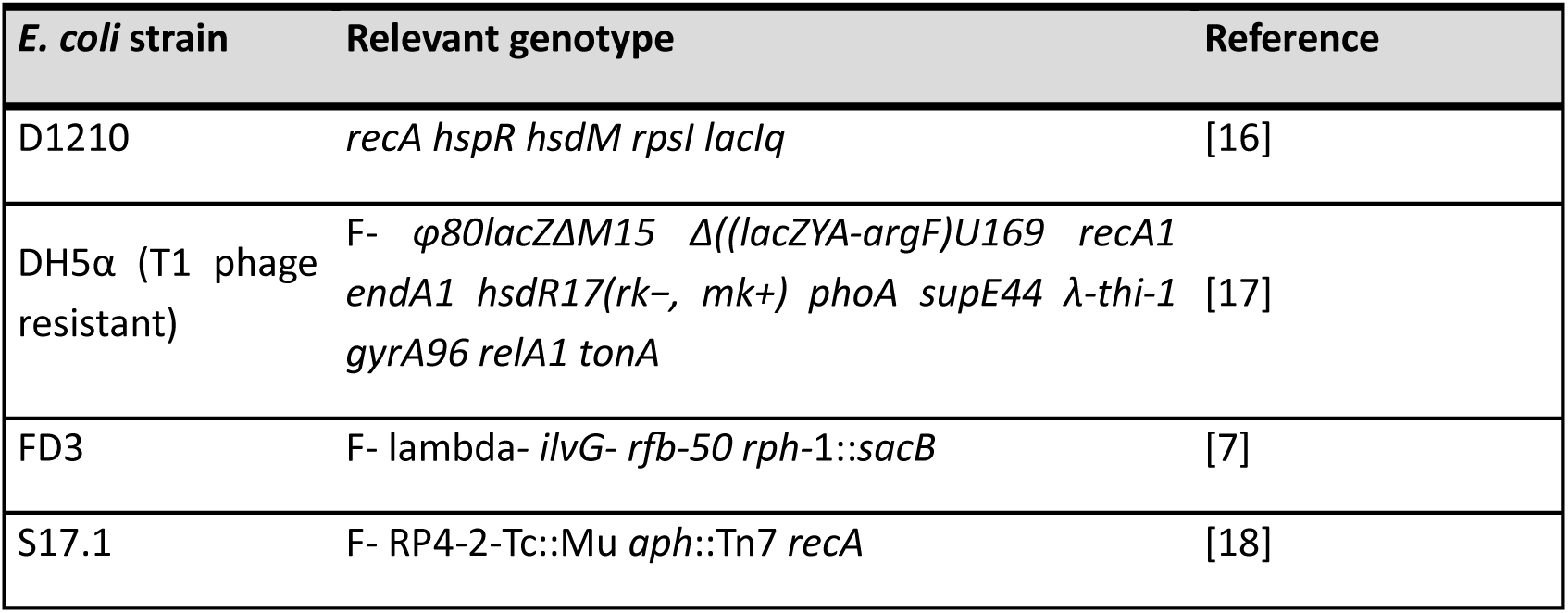
Escherichia coli strains.

| ***E. coli*** **strain** | **Relevant genotype** | **Reference** |
|---|---|---|
| D1210 | *recA hspR hsdM rpsI lacIq* | [16] |
| DH5α (T1 phage resistant) | F- *φ80lacZΔM15 Δ((lacZYA-argF)U169 recA1 endA1 hsdR17(rk−, mk+) phoA supE44 λ-thi-1 gyrA96 relA1 tonA* | [17] |
| FD3 | F- lambda- *ilvG- rfb-50 rph*-1::*sacB* | [7] |
| S17.1 | F- RP4-2-Tc::Mu *aph*::Tn7 *recA* | [18] |

### 2.2. Plasmid used and constructed in this work

Plasmids used in this work (Table 2) were constructed by Gibson assembly [19]. Backbones and inserts were amplified by PCR using PCRBIO HiFi polymerase, with insert primers carrying 40-nt overhangs homologous to the adjacent fragments. For pHOT06 and pHOT07, annealed oligonucleotides encoding the gRNA spacer were ligated into the BsaI-digested vector pHOT04. Oligonucleotides were designed with SnapGene, targeting a Tm of approximately 60 °C, and were synthesized by Sigma-Aldrich (Merck KGaA, Darmstadt, Germany). Oligonucleotides used for plasmid construction and verification are listed in Table 3.

**Table 2:**
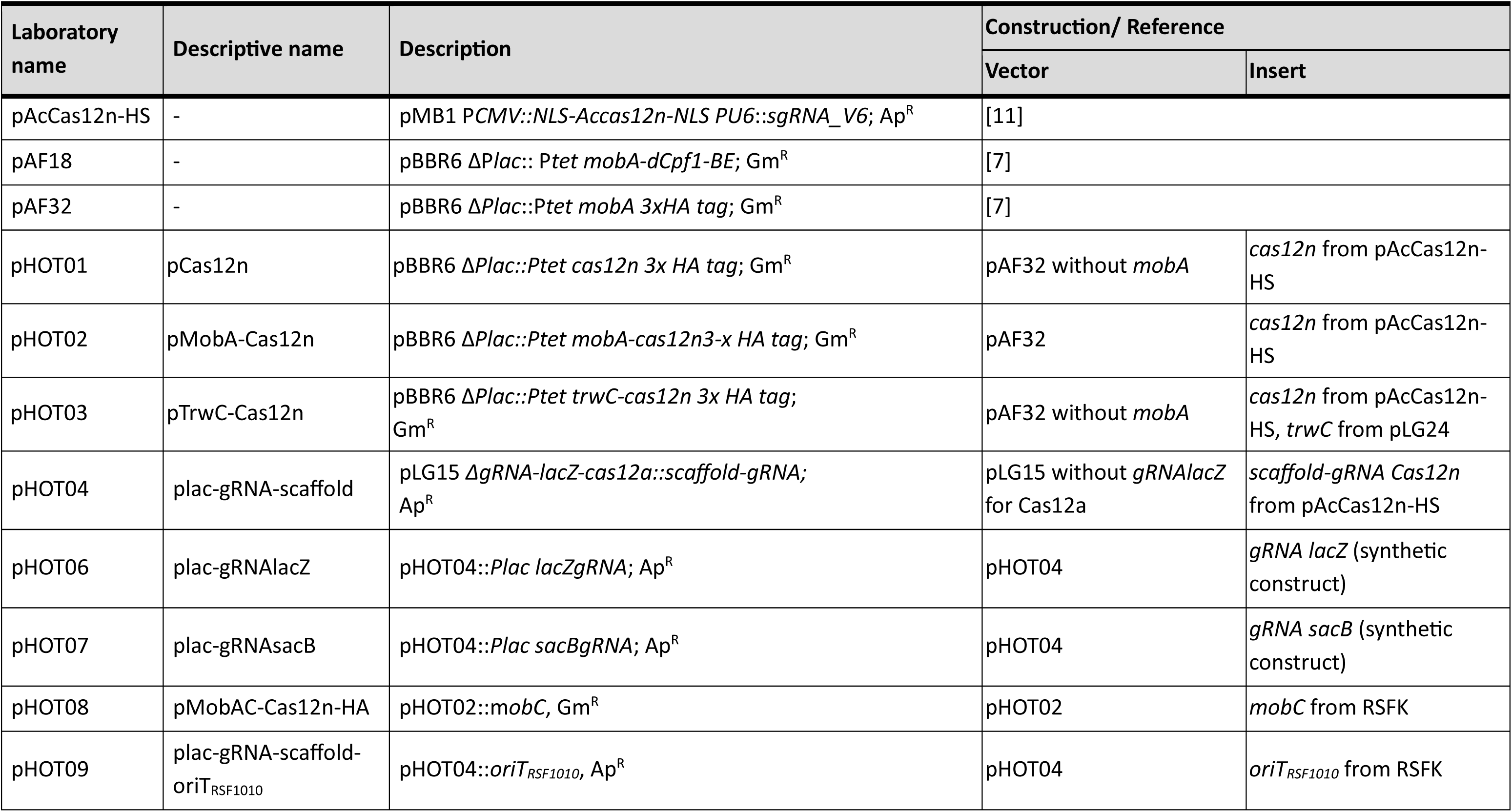
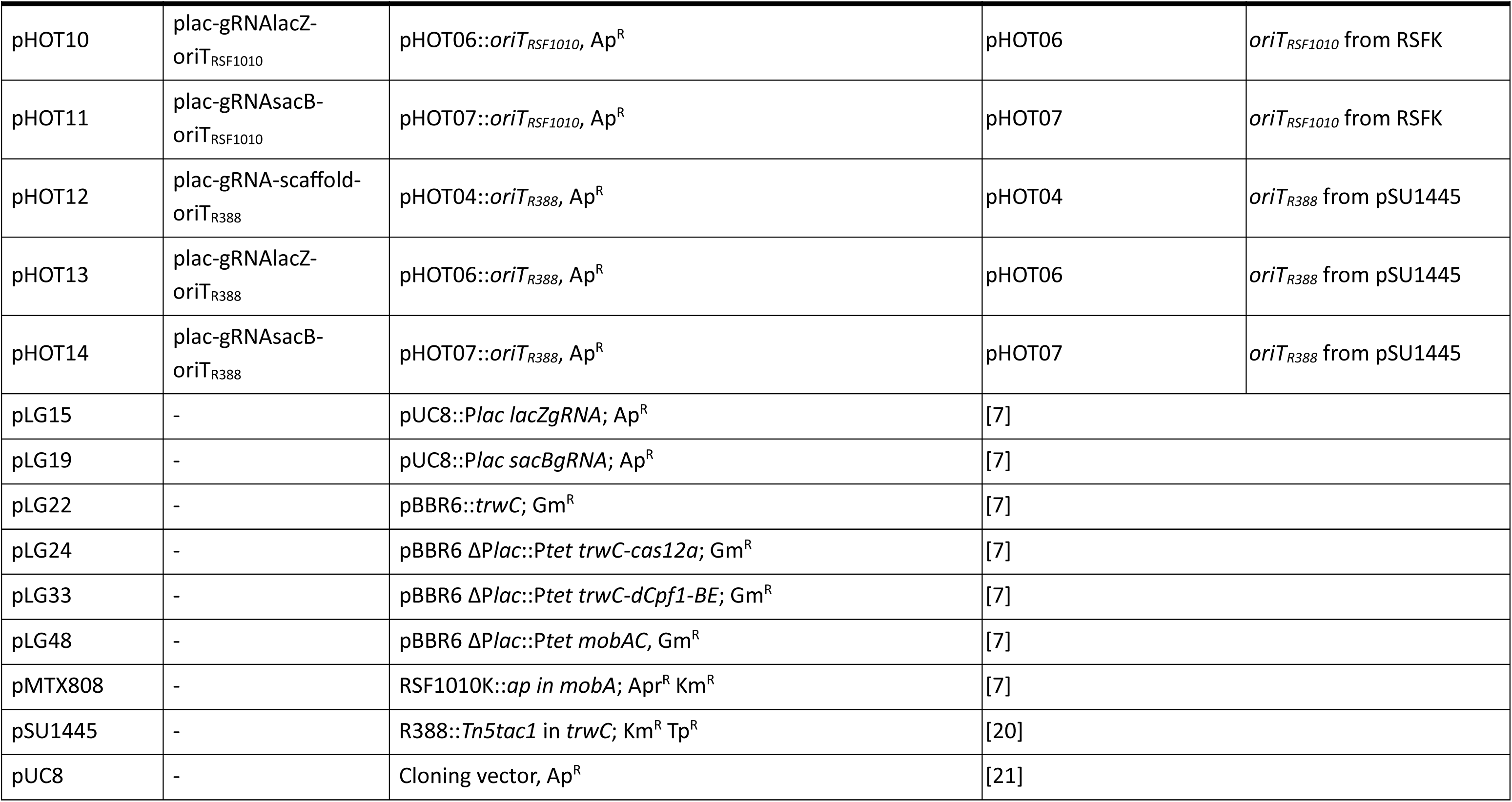
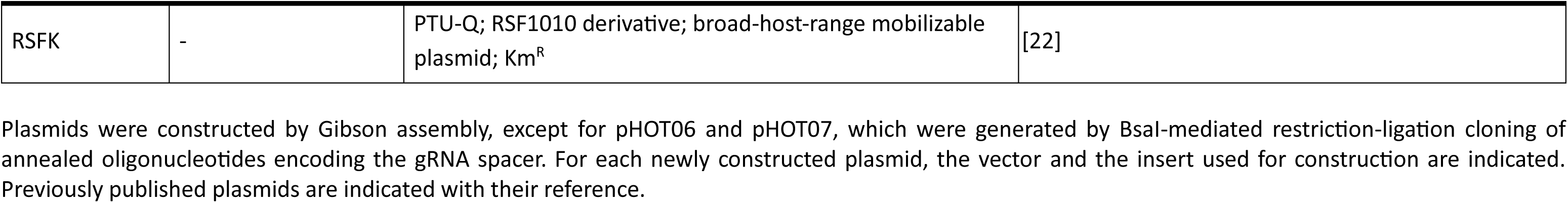
Plasmids used in this work.

**Table 3:**
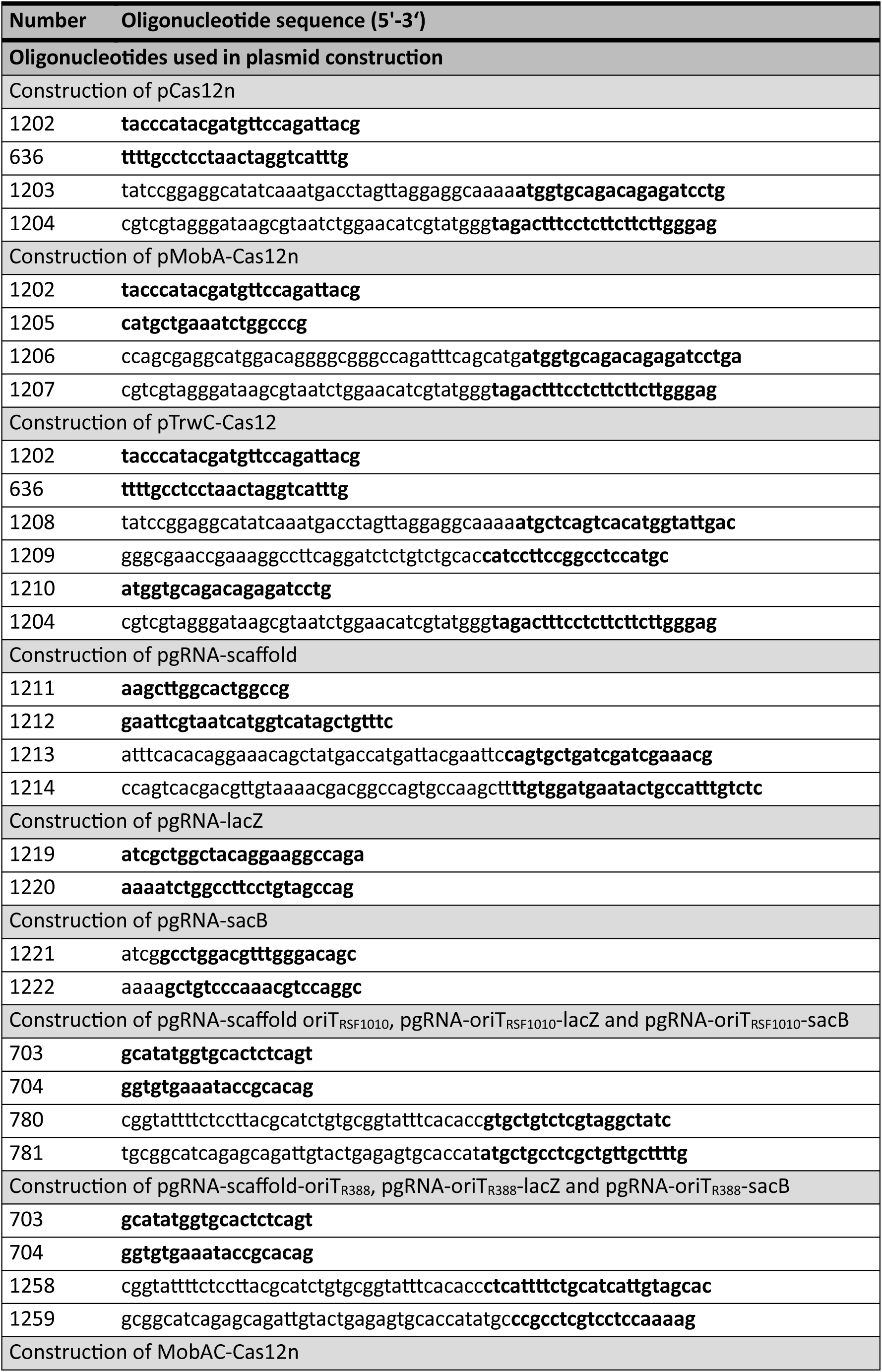
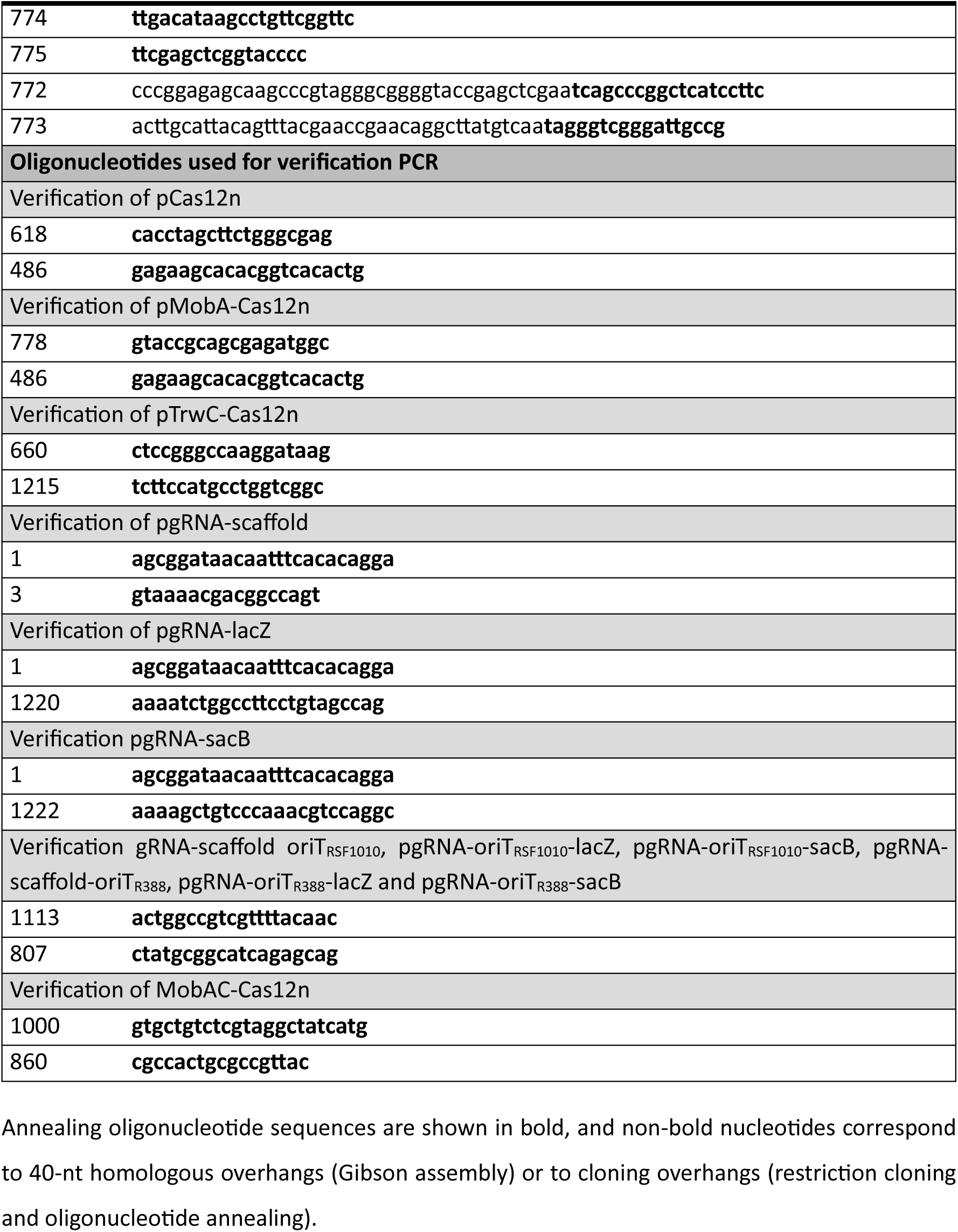
Oligonucleotides used in this work.

Annealing oligonucleotide sequences are shown in bold, and non-bold nucleotides correspond to 40-nt homologous overhangs (Gibson assembly) or to cloning overhangs (restriction cloning and oligonucleotide annealing).

### 2.3. Western blot analysis

Cultures of bacteria harboring the respective plasmids were grown overnight and diluted 1:20 in fresh LB. Protein expression was induced for 3 h with 500 µM IPTG or 200 ng/ml aTc, according to the plasmid. For each culture, 1 mL was harvested by centrifugation, and the pellet was resuspended in 100 µL of 2× SDS-gel loading buffer. Lysates were kept at −20 °C for at least overnight, and loading volumes were normalized to the OD600 of each culture. After boiling for 5 min, samples were separated on 9% sodium dodecyl-sulfate polyacrylamide gel electrophoresis (SDS-PAGE) using Tris-glycine-SDS running buffer. Proteins were subsequently transferred to nitrocellulose membranes. Membranes were then probed with one of the primary antibodies anti-TrwC [23] (1:10,000) or anti-HA-tag rabbit polyclonal (Proteintech; 1:10,000), and with the secondary antibody anti-rabbit IgG IRDye® 800CW (LI-COR; 1:10,000). Signals were detected using an Odyssey CLx imaging system (LI-COR), and NZYColour Protein Marker II (NZYTech) served as molecular weight standard.

### 2.4. Mating Assays

Mating assays were performed on solid LB medium in 24-well plates. Donor and recipient strains were grown overnight in LB medium supplemented with the appropriate antibiotics and diluted 1:20 in fresh medium containing the relevant inducers (IPTG and/or aTc). Cultures were incubated for an additional 3 h before cells were harvested by centrifugation (3.380 × g, 5 min), resuspended in fresh LB medium, and mixed at a 1:1 ratio based on OD_600_. The mixtures were centrifuged again, resuspended in 15 µL LB medium, and spotted onto solid LB medium supplemented with the appropriate inducers (IPTG and/or aTc). After drying, mating plates were incubated at 37 °C for 1 or 3 h, depending on the experiment. Following mating, serial dilutions were plated on selective LB agar to enumerate donors, recipients, and transconjugants. Plates were incubated overnight at 37 °C, and colony-forming units (CFUs) were determined the following day. Conjugation frequencies were calculated as the ratio of transconjugants to donor (T/D) or recipient (T/R) cells, as indicated.

#### 2.4.1. Conjugation complementation assays

Conjugation complementation assays were performed to assess whether the relaxase domain within the Relaxase-Cas12n fusion proteins retains its activity. The assay relies on complementation in trans of a mobilizable plasmid carrying a nonfunctional relaxase gene to restore plasmid mobilization. For the RSF1010 system, *E. coli* S17.1 carrying pMTX808, which provides the RP4 conjugative machinery and the components required for MobA-mediated mobilization but lacks a functional *mobA* gene, was used as the donor strain. Donor cells were complemented with either pMobA-Cas12n or pAF18, encoding the MobA-Cas12n fusion protein or wild-type MobA, respectively. For the R388 system, *E. coli* D1210 carrying pSU1445, which provides the components required for TrwC-mediated mobilization but lacks a functional *trwC* gene, was used as the donor strain. These cells were complemented with either pTrwC-Cas12n or pLG22, encoding the TrwC-Cas12n fusion protein or wild-type TrwC, respectively. *E. coli* DH5α was used as the recipient strain in both systems. Mating assays were performed as described above, with a mating time of 1h.

### 2.5. Lethality assays

Lethality assays were performed to assess the functionality of the Cas domain within the Relaxase-Cas fusion proteins. *E. coli* D1210 carrying pMobA-Cas12n, pTrwC-Cas12n or pCas12n were used to prepare electrocompetent cells, which were transformed with 100 ng of pgRNA-lacZ or pgRNA-sacB. Following transformation, serial dilutions were plated on LB agar supplemented with appropriate antibiotics with or without aTc and IPTG to induce expression of the fusion proteins and gRNA, respectively. When non-specific cleavage was observed upon induction of MobA-Cas12n, the induction period was shortened as follows. Following electroporation and a 1 h recovery period, an aliquot of the transformation mixture was removed and plated in serial dilutions on non-inducing medium. Inducers were then added to the remaining transformation mixture, and cells were incubated for an additional 2 h before being plated in serial dilutions on non-inducing medium. Plates were incubated overnight at 37 °C, and CFUs were quantified to assess Cas-mediated lethality under the different induction conditions.

#### 2.5.1. Lethality assays after translocation

Lethality assays in recipient bacteria after Relaxase-Cas translocation were performed to assess whether the Cas domain regained its nuclease activity following translocation into recipient cells. Donor strains (*E. coli* S17.1 or D1210 for the relaxases MobA and TrwC, respectively) carried i) the components required for conjugation, ii) non-mobilizable plasmids encoding the Relaxase-Cas fusion proteins (pMobA-Cas12n or pTrwC-Cas12n), or relaxase alone as controls, and iii) mobilizable gRNA-encoding plasmids. For the RSF1010 system, donors carried either plac-gRNAsacB-oriT_RSF1010_, encoding a *sacB*-targeting gRNA, or plac-gRNA-scaffold-oriT_RSF1010_, encoding the scaffold gRNA as a negative control. For the relaxase TrwC, the corresponding plasmids were plac-gRNAsacB-oriT_R388_ and plac-gRNA-scaffold-oriT_R388_, respectively. Conjugation was performed as described above, with a mating time of 3 h.

### 2.6. Statistical Analysis

Statistical significance was assessed using parametric or non-parametric tests, as appropriate based on normality and homogeneity of variance. One-way ANOVA or unpaired Student’s *t*-tests were used for parametric data, whereas Kruskal-Wallis or Mann-Whitney *U* tests were applied to non-parametric data. The statistical test used for each comparison is specified in the corresponding figure legend. All experiments were performed with at least three independent biological replicates (n ≥ 3), and statistical analyses were conducted using GraphPad Prism 8.

## 3. Results

### 3.1. Production and stability of the Relaxase-Cas fusion proteins

To test whether a miniature nuclease can be delivered by conjugation to the recipient bacteria, we constructed two fusion proteins in which Cas12n was fused to the C-terminus of a conjugative relaxase: MobA from the mobilizable system RSF1010 and TrwC from conjugative system R388. We added a 3×HA tag at the C-terminus of each construct to facilitate western blot detection. We also generated a plasmid expressing Cas12n alone as a nuclease-only control. The expression of the fusions was controlled by the P*tet* promoter.

We examined the production and stability of the fusions by western blot on total protein extracts of donor cells (Fig 2). We could observe proteins of the predicted sizes: 139 kDa for MobA-Cas12n and 168 kDa for TrwC-Cas12n, compared with 263 kDa for the TrwC-Cas12a fusion of the original ConCasTool. We used both anti-HA and anti-TrwC antibodies (when applicable) for detection, since the contrast between the two antibodies is informative: fragments that retain TrwC epitopes but have lost the C-terminal HA tag are consistent with proteolysis occurring within Cas12n or close to the C terminus of the fusion. First, MobA-Cas12n and TrwC-Cas12n were both detected by using an anti-HA antibody (Fig 2A). A predominant band at their expected sizes were detected. We also detected, especially for TrwC-Cas12n, minor lower-molecular-weight species, probably due to protein degradation (Fig 2). The anti-TrwC antibody allowed us to compare the new TrwC fusion with the previously constructed fusion TrwC-Cas12a (Fig 2B). Both proteins gave numerous lower-molecular-weight species. In addition, we could observe the full-length protein detectable only as a faint band in the case of TrwC-Cas12a. Thus, under these conditions, the Cas12n fusions appeared more stable than TrwC-Cas12a, and MobA-Cas12n seemed less susceptible to proteolysis. Both factors may increase the amount of intact protein available for translocation.

**Fig 2.**
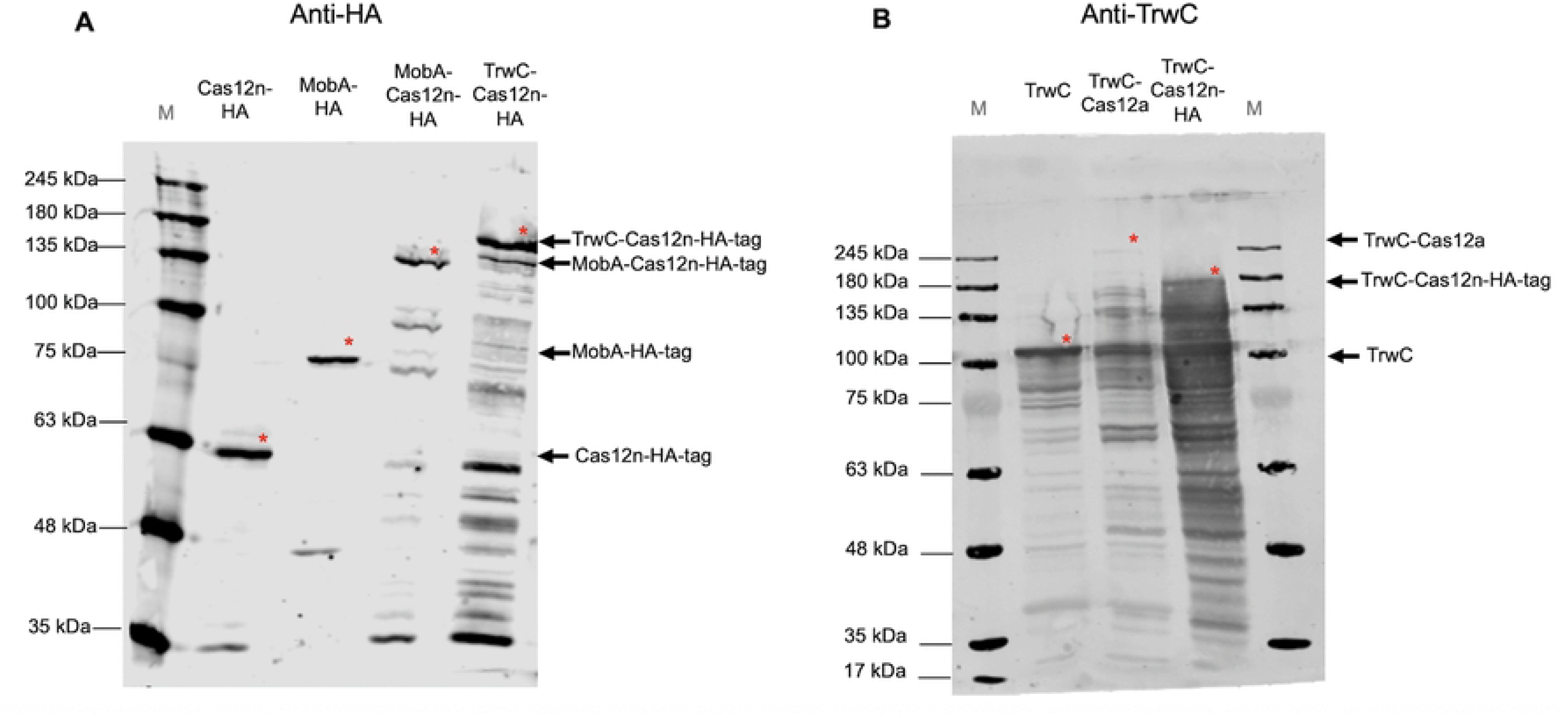
Production and stability of Cas12n fusion proteins. Samples correspond to induced *E. coli* D1210 cell lysates carrying plasmids which produce the proteins indicated at the top. M, NZY Colour Protein Marker II (sizes indicated to the left. A) Western blot analysis using an anti-HA-tag primary antibody. B) Western blot analysis using an anti-TrwC antibody. Black arrows point to the expected migration of the indicated proteins, and red asterisks indicate the corresponding observed bands. Expected sizes are: Cas12n-HA-tag: 61 kDa, MobA-Cas12n-HA 139 kDa, TrwC-Cas12n-HA 168 kDa, MobA-HA 81 kDa, TrwC 107 kDa, TrwC-Cas12a 263 kDa.

### 3.2. The fusion proteins retain relaxase and site-specific nuclease activity

To confirm that the fusion proteins were functional and that the activity of each domain is not affected in the fusion, we assayed the site-specific nuclease and the relaxase activity within the donor bacteria. Because double-strand-breaks (DSBs) are generally lethal in bacteria, which lack efficient non-homologous end joining, Cas12n site-specific nuclease activity can be assayed as a loss of viability when the nuclease is directed against a chromosomal gene. As target for the assay, we selected the chromosomal genes *lacZ* (present in the chromosome) and *sacB* as negative control, since it is not present in the chromosome (Fig 3A); both targets were previously used in the characterization of TrwC-Cas12a [7]. gRNAs were expressed under the P*lac* promoter from the scaffold optimized for Cas12n [11], and target sequences were introduced into the scaffold (see Methods). Donor cells carrying each fusion construct were electroporated with a plasmid encoding gRNAs targeting either *lacZ* or *sacB* and plated with or without inducers. The results are shown in Fig 3B and 3C. For TrwC-Cas12n and for Cas12n alone, we obtained the expected result: when the cell was electroporated with the *lacZ-*gRNA under induction conditions, a reduction in the number of colonies of approximately four to five orders of magnitude was obtained when compared to cells receiving the non-targeting guide. Also, no reduction was observed in the absence of inducers, confirming both guide-dependent cleavage and tight control of expression. However, MobA-Cas12n fusion protein behaved differently. Induction of its expression reduced colony numbers, whether or not the gRNA matched a chromosomal target, which indicated an effect independent of target recognition, and suggested that overproduction of this fusion was itself toxic (Fig 3B). In order to fine tune the activity of the fusion and to decrease the possible toxicity effect, we reduced the induction times from overnight to 2 h, after the electroporation and before plating the mixture (see Methods). The results are shown in Table 4. Under these conditions, a reduction of approximately one order of magnitude was observed only when both the *lacZ* guide and the inducers were present, and not with the non-targeting guide. These results showed that MobA-Cas12n retains target-specific nuclease activity, accompanied by a toxicity that is independent of target recognition and that can be managed by limiting its expression.

**Fig 3.**
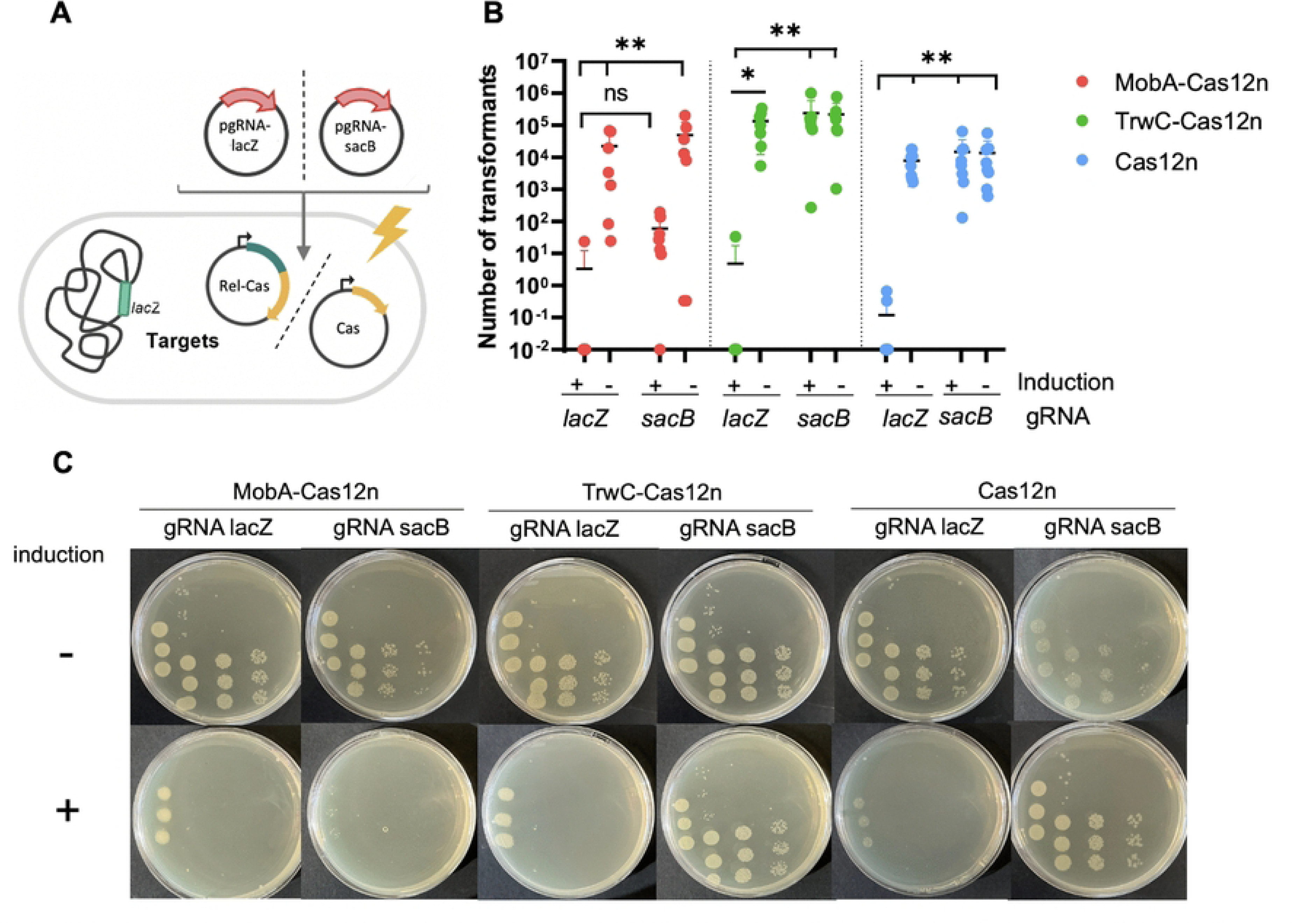
Site-specific endonuclease activity of Cas12n fusion proteins. A) Schematic overview of the assay. *E. coli* D1210 harboring the plasmids which produce the indicated proteins were transformed with pgRNA-lacZ or pgRNA-sacB. B) Number of transformants after electroporation of plasmids encoding gRNAs into *E. coli* D1210 carrying plasmids encoding Cas12n and Cas12n-relaxase fusions. Statistical analysis was performed using a Kruskal-Wallis test (*p < 0.05, **p < 0.01, ***p < 0.001). C) Representative agar plates displaying the number of colonies upon transformation and induction (+) vs. non-induction (-). Induction of the expression of the fusion proteins and gRNAs with aTc and IPTG respectively is indicated as “+” and compared to non-induction.

**Table 4:** Lethality assay for MobA-Cas12n with short induction times.

| **Transformed plasmid** | **No. of CFU** |  |
|---|---|---|
|  | **No induction** | **2 h induction** |
| **pgRNA-lacZ** | 1,13E+04 ± 1,13E+04 | 4,62E+03 ± 6,19E+03 |
| **pgRNA-sacB** | 5,15E+04 ± 5,44E+04 | 2,02E+04 ± 1,39E+04 |

*E. coli* D1210 carrying pMobA-Cas12n was electroporated with plasmids encoding gRNA for *lacZ* or *sacB,* and incubated under conditions of non-induction or induction with IPTG and aTc for 2h (n=2). The table shows the number of transformants obtained (mean +/- standard deviation).

The relaxase activity of the fusion proteins was analyzed by conjugation complementation assays, in which a helper plasmid carrying a disrupted relaxase gene is mobilized by a functional relaxase supplied in trans. We constructed donor cells carrying the systems for mobilization of the corresponding relaxases (Fig 4A): for translocation of TrwC fusions, a plasmid R388 derivative deficient in *trwC*; for translocation of MobA fusions, a plasmid RSF1010 derivative deficient in *mobA*, plus the chromosomally-encoded RP4 system to provide the T4SS. The results of the conjugation assays are shown in Fig 4B. In both systems, the fusion proteins mobilized the helper plasmid, with no significant difference from the corresponding wild-type relaxase. It is also observed that the conjugation frequencies are higher for the MobA complementation assays; however, this is likely due to the above-mentioned toxicity of MobA overexpression, which leads to lower number of viable donor cells, affecting the frequencies expressed as transconjugants per donor cell.

**Fig 4.**
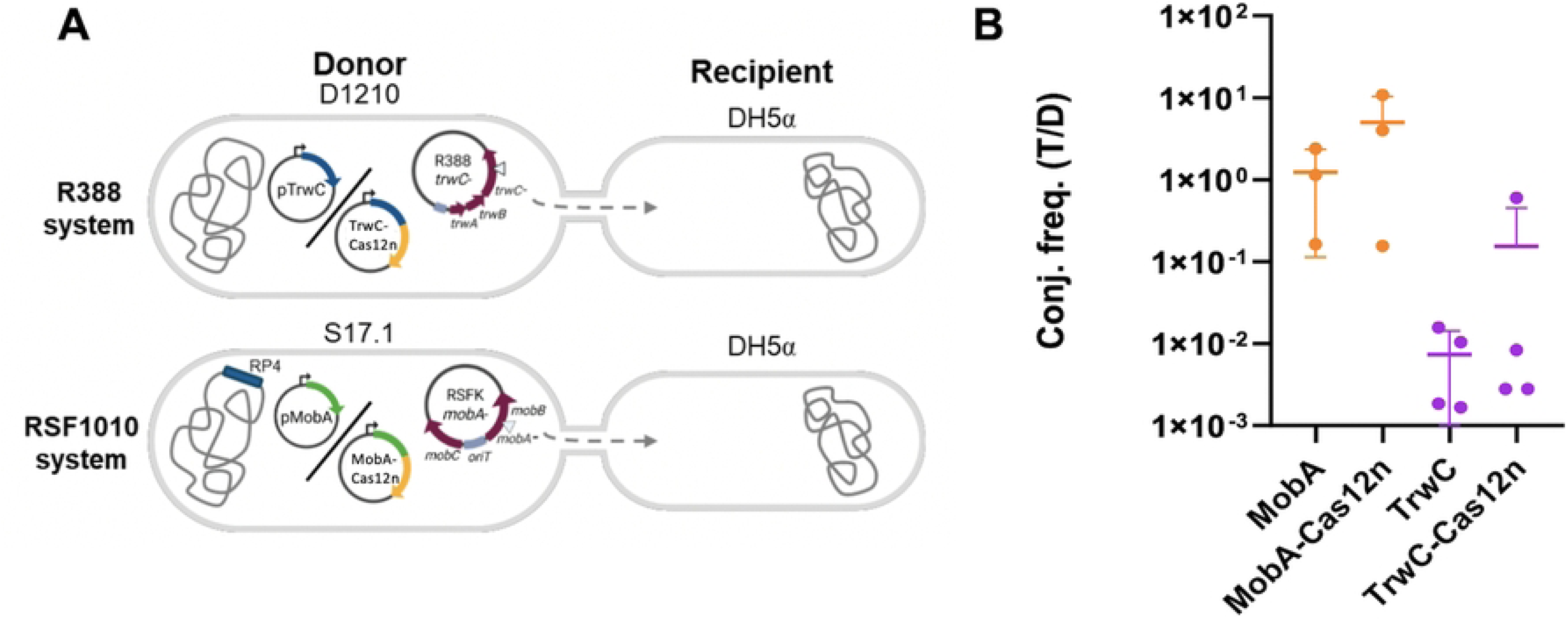
Assessment of the relaxase activity by complementation assays. A) Schematic representation of the complementation assays. In both the R388 system (top) and the RSF1010 system (bottom), helper plasmids with non-functional relaxase genes are complemented in trans either by the WT relaxase or by the corresponding Relaxase-Cas12n fusions. In the case of RSF1010, RP4 inserted in the chromosome is also required, to provide the T4SS. B) Conjugation frequencies of relaxase-deficient mobilizable plasmids when complemented with fusion proteins MobA-Cas12n and TrwC-Cas12n compared to those obtained using the corresponding wild-type relaxases, MobA and TrwC. Conjugation frequencies were calculated a transconjugant per donor (T/D). No statistically significant differences could be found using a two-tailed unpaired Student’s t-test for MobA/MobA-Cas12n comparisons and a Mann-Whitney U test for TrwC/TrwC-Cas12n comparisons (*p < 0.05, **p < 0.01, ***p < 0.001).

### 3.3. Evidence for Cas12n activity in the recipient upon translocation during conjugation

Previous work with TrwC-Cas12a successfully used a lethality assay to measure RNA-guided nuclease activity in the recipient upon targeting a *sacB*-containing recipient with anti-*sacB* gRNA [7]. Moreover, we showed that the assay is more efficient when the gRNA is delivered from the donor on a mobilizable plasmid, co-translocated covalently attached to the fusion protein, than when it is supplied by prior transformation of the recipient [7]. This configuration also has the practical advantage of requiring no modification of the recipient. We therefore applied the same strategy to the Cas12n fusions.

To this end, plasmids encoding the gRNA had to became mobilizable from donor to recipient bacteria, so the *oriT* of RSF1010 or of R388 was added to each guide plasmid. Also, to reduce the number of plasmids residing in the donor cells, we included the *mobC* gene into pMobA-Cas12n so that the plasmid contains all the elements needed for DNA recruitment without the need to add an RSF1010 derivative. Fig 5A summarized the experimental setup. We generated donors carrying the fusion protein or the wild-type relaxase together with a mobilizable plasmid encoding either a *sacB*-targeting gRNA or only the gRNA scaffold. As recipient, we used *E. coli* carrying a chromosomal copy of *sacB*. TrwC-Cas12a was assayed in parallel under the same conditions.

**Fig 5.**
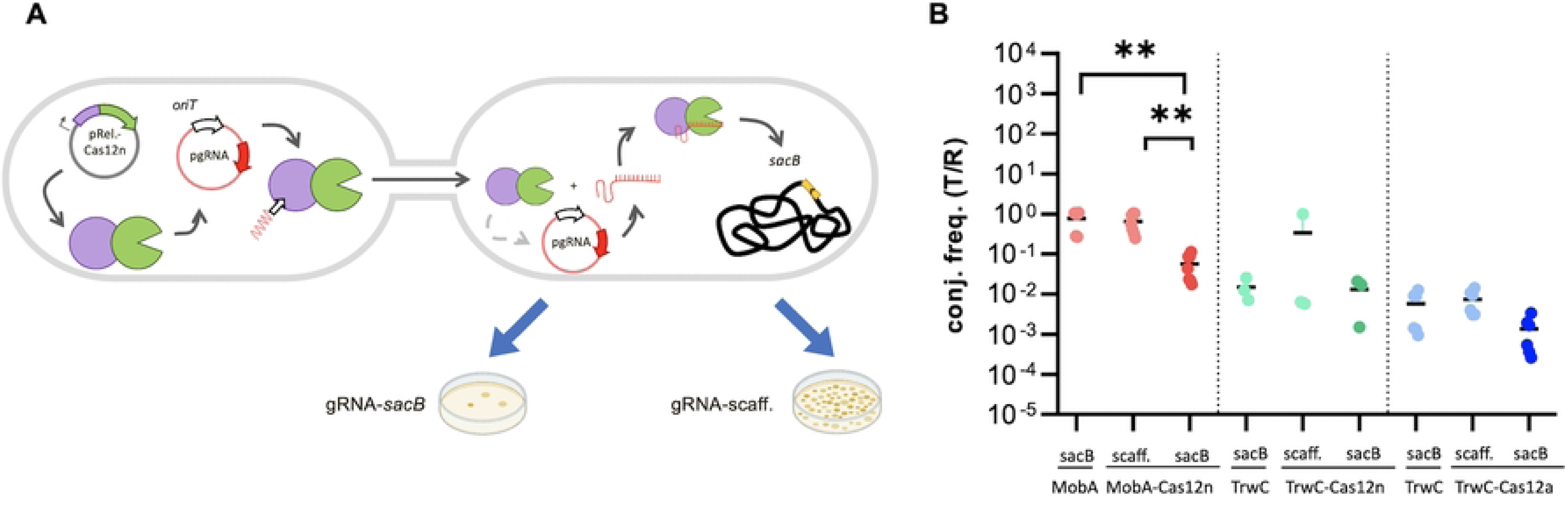
Lethality assay in transconjugants following translocation of Relaxase-Cas fusions. A) Schematic overview. Donor cells carry a Relaxase-Cas fusion and a mobilizable plasmid encoding either a *sacB*-targeting gRNA (*sacB*) or a scaffold control (*scaff.*). Following nicking at *oriT*, the relaxase-DNA complex is transferred via the T4SS into FD3 recipient cells, where the gRNA directs Cas to chromosomal *sacB*. Target cleavage is expected to reduce recipient viability and transconjugant recovery compared with the scaffold gRNA control. B) Conjugation frequencies (T/R) for MobA-Cas12n, TrwC-Cas12n, and TrwC-Cas12a, compared with the respective wild-type relaxase controls. Statistical significance was assessed using the Kruskal-Wallis test (MobA-Cas12n and TrwC-Cas12n) or one-way ANOVA (TrwC-Cas12a) (*p < 0.05, **p < 0.01, ***p < 0.001).

For MobA-Cas12n, when the *sacB-*gRNA was translocated to the recipient bacteria, a decrease in the number of transconjugants of more than one order of magnitude was observed. This reduction was not observed in the absence of the *sacB*-specific gRNA, confirming that MobA-Cas12n was translocated to the recipient cell, where it regained its target-specific activity. By contrast, the slight decrease observed for TrwC-Cas12n and TrwC-Cas12a was not statistically significant. So, in addition to showing that MobA-Cas12n can be translocated and active in recipient cells, our data suggest that this new fusion protein is more efficient than TrwC-Cas12a tested in parallel.

## 4. Discussion

The ability to genetically modify bacteria is essential for a wide range of biotechnological applications, but efficient delivery of genome-editing tools such as CRISPR-Cas systems remains a major challenge, particularly for wild-type bacteria, which are often considerably harder to transform than laboratory strains like *E. coli*. ConCasTool addresses this problem by delivering Cas proteins, fused to conjugative relaxases, directly to recipient cells via bacterial conjugation, circumventing the need for CRISPR-Cas expression in the recipient cell itself [7]. Nevertheless, the current system achieves limited editing efficiency, with only 1 in 50,000 transconjugants being successfully edited, even when a HR template is provided and *E. coli* is used as the recipient strain [7]. Given that wild-type bacteria are inherently more recalcitrant to genetic modification [24] and often exhibit lower conjugation frequencies than *E. coli* laboratory strains [15], improving the overall efficiency of this system is a prerequisite for its broader applicability. To improve ConCas efficiency, we hypothesized that increasing Relaxase-Cas fusion stability could be beneficial. We reasoned that alternative Cas variants could improve fusion stability, minimize proteolysis, and facilitate post-translocation refolding. For these reasons, we selected the alternative nuclease Cas12n [11] in place of Cas12a. Another goal of this work was to expand the range of potential recipient cells for ConCasTool delivery, for which the promiscuous relaxase MobA was included in this study.

We engineered fusions of the conjugative relaxases MobA and TrwC to Cas12n, as well as plasmids expressing gRNAs designed against *lacZ* and *sacB*. Western blot analysis confirmed expression of all constructs under the P*tet* promoter, with bands at the expected molecular weights. Degradation was particularly pronounced for TrwC-Cas12n. Analysis using an anti-HA antibody detected mainly the full-length protein, whereas anti-TrwC revealed numerous shorter species, indicating that cleavage occurs within Cas12n or close to the C-terminal tag. Anti-HA Western blot analysis revealed comparable expression levels of MobA-Cas12n and TrwC-Cas12n. Comparison of TrwC-Cas12n and TrwC-Cas12a in the anti-TrwC Western blot further showed a markedly stronger full-length signal for TrwC-Cas12n relative to TrwC-Cas12a, the latter being detectable only as a faint band with extensive degradation. This degradation was markedly reduced in the case of MobA-Cas12n. Taken together, these results indicate that the Cas12n-based fusions may increase the amount of intact fusion protein available for T4SS translocation.

Previous studies had shown that both MobA and TrwC remain functional when fused to Cas12a or base editors in the ConCas system. However, as Cas12n is a distinct and recently characterized Cas protein, the functionality of both the relaxase and endonuclease domains had to be experimentally validated in the newly generated fusion proteins. Relaxase functionality was evaluated in conjugation complementation assays. Both fusion proteins successfully mobilized a relaxase-deficient helper plasmid, suggesting that the relaxase domain retains its activity when incorporated into a fusion protein. Similar to previous results, the fusion proteins seemed to be toxic to donor cells, as evidenced by a reduction in viable donors. On the other hand, Cas12n activity was assessed by transformation-based lethality assays. For TrwC-Cas12n and the Cas12n control, induction in cells transformed with the target gRNA plasmid pgRNA-lacZ resulted in a drastic reduction in CFU compared to cells carrying the non-target pgRNA-sacB. These results demonstrate that the Cas12n domain retains target-specific DNA cleavage activity when expressed as part of a fusion protein specifically under induction conditions. In contrast, induction of MobA-Cas12n resulted in a comparable reduction in CFU under the same conditions regardless of whether cells carried the target or non-target gRNA plasmid, indicating non-specific cleavage activity. Attributing this to toxicity of the overexpressed fusion, we limited the induction time, after which colony numbers fell only when both the *lacZ* gRNA and the inducer were present, and not with the *sacB* gRNA or in the absence of induction, thus confirming the site-specific nuclease activity of the fusion protein.

Toxicity of proteins with DNA binding and cleaving activities, such as relaxases and Cas nucleases, is a common observation when overexpressed, and it can represent a serious obstacle for experimental design. In this case, however, since our goal is to test the activity of the Cas proteins upon translocation into recipient cells, where a limited number of protein molecules are translocated and recover their function, toxicity is not a concern; in fact, this is the basis of our rationale for delivering Cas proteins instead of *cas*-encoding DNA.

The substantial degradation observed by Western blot analysis, particularly for TrwC-Cas12n, raises the possibility that free relaxase and/or Cas12n domain, released through proteolytic cleavage of the fusion protein, may retain their respective activities. Consequently, the above results alone do not constitute sufficient evidence for transfer of the intact fusion protein. Assessing Cas12n endonuclease activity in recipient cells provides a more stringent readout, since detecting nuclease activity requires not only translocation but also correct refolding of the protein after the at least partial unfolding known to occur during T4SS translocation [8,9]. Relaxase-Cas fusions with MobA and TrwC have previously shown to remain functional after translocation for both Cas12a and base editors, and functional delivery of CRISPR-associated proteins via T4SS has also been reported in other systems, such as VirF-Cas9 fusions delivered to plant cells through the *Agrobacterium* VirB/D4 system [25]; however, these findings do not necessarily predict the behavior of Cas12n. Experimental validation was therefore required to assess whether Relaxase-Cas12n fusions can be translocated through the T4SS and are able to refold and recover endonuclease activity in recipient cells.

Our group had previously demonstrated the highest editing efficiency of ConCasTool when the gRNA-encoding plasmid was delivered from the donor to the recipient as a mobilizable plasmid co-translocated with the fusion protein. This improvement was proposed to result from coordinated delivery of both components, increasing the likelihood of productive Cas-gRNA complex formation immediately after transfer. Thus, we applied this strategy to Cas12n-mediated gene editing. Analysis of conjugation frequencies after fusions translocation revealed a significant reduction of more than >1-log for donors carrying pgRNA-*sacB*-*oriT*_RSF1010_ compared to donors carrying control plasmids with no targeting gRNA. No comparable reduction was observed for TrwC-Cas12n under the same conditions. Our results provide evidence that the MobA-Cas12n fusion protein retains Cas12n nuclease activity after translocation into recipient cells.

The values previously reported for TrwC-Cas12a were calculated using conjugation frequencies expressed as transconjugants per donor; in these experiments, however, the toxicity of the fusions and the consequent loss of donor viability made it necessary to express the data as transconjugants per recipient, so the absolute frequencies obtained in the two studies cannot be compared directly. To allow a comparison between the activity of the fusions, experiments including TrwC-Cas12a were performed in parallel; gene targeting efficiency, as inferred from the lethality assays, was higher for MobA-Cas12n than for TrwC-Cas12a, although the reduction obtained for TrwC-Cas12a was itself smaller than that previously reported. Finally, it is noteful that we did not detect TrwC-Cas12n activity in recipient cells, even though both moieties of the fusion were functional in other constructs (TrwC-Cas12a and MobA-Cas12n). This demonstrates that each fusion protein has to be tested individually for activity in the recipient cell.

The higher expression levels observed for the Cas12n fusions may help overcome one of the main limitations of TrwC-Cas12a, namely the low amount of fusion protein detected. Moreover, the use of MobA as the conjugative relaxase expands the potential host range of the system, as the RSF1010 mobilization machinery is recognized by a broad variety of conjugative T4SS and especially by RP4, whose promiscuity goes beyond bacteria. This could enable delivery to recipient species that are not accessible through the R388 system, thereby broadening the applicability of the ConCas platform. Taken together, these results demonstrate that the newly generated MobA-Cas12n fusion constitutes a successful and effective alternative to the previously developed TrwC-Cas12a system.

The development of a conjugation-based CRISPR delivery system that circumvents the need to express the Cas nuclease in recipients, and that is capable of reaching a broad host-range of wild-type bacteria, represents a significant advance in the field of bacterial genome engineering. Conventional genetic manipulation relies on transformation-based approaches, which are inherently limited to subset of naturally competent or laboratory adapted strains. Conjugation, by contrast, is a near-universal mechanism of horizontal gene transfer that operates across a wide diversity of Gram-negative and Gram-positive bacteria, including species that remain inaccessible using standard molecular techniques. A functional ConCasTool, which does not require fine-tuned expression of the Cas nuclease in the recipient, would therefore provide access to bacterial species that are currently inaccessible to standard genetic manipulation, including environmentally, industrially, and clinically relevant microorganisms. The demonstration that MobA-Cas12n is properly expressed and that it successfully retains nuclease activity in recipient cells following translocation, represents a substantial and concrete step toward this goal.

## 5. Acknowledgements

This work was funded by grant PID2023-147192OB-I00 from MICIU/AEI/10.13039/501100011033/ and ERDF/EU to ML. JH was a recipient of an Erasmus+ traineeship grant from the European Union and a JAE Intro ICU fellowship (JAEICU_25_03779) from the Spanish Research Council (Consejo Superior de Investigaciones Científicas, CSIC). We are grateful to Andrea Fernández-Gómez for help with the Figures, and to Ivana González for technical support.

## Notes

### Competing Interest Statement

The authors have declared no competing interest.

## References

1. Yan X, He Q, Geng B, Yang S. Microbial cell factories in the bioeconomy era: from discovery to creation. BioDesign Res. 2024;6: 0052.

2. Pacesa M, Pelea O, Jinek M. Past, present, and future of CRISPR genome editing technologies. Cell. 2024;187: 1076–1100.

3. Zalila-Kolsi I. Engineered bacteria as living therapeutics: Next-generation precision tools for health, industry, environment, and agriculture. AIMS Microbiol. 2025;11: 946.

4. Cui L, Vigouroux A, Rousset F, Varet H, Khanna V, Bikard D. A CRISPRi screen in *E. coli* reveals sequence-specific toxicity of dCas9. Nat Commun. 2018;9: 1912.

5. Lu L, Shen X, Sun X, Yan Y, Wang J, Yuan Q. CRISPR-based metabolic engineering in non-model microorganisms. Curr Opin Biotechnol. 2022;75: 102698.

6. Garcillán-Barcia MP, de la Cruz F, Rocha EPC. The extended mobility of plasmids. Nucleic Acids Res. 2025;53: gkaf652.

7. Guzmán-Herrador DL, Fernández-Gómez A, Depardieu F, Bikard D, Llosa M. Delivery of functional Cas:DNA nucleoprotein complexes into recipient bacteria through a type IV secretion system. Proc Natl Acad Sci. 2024;121: e2408509121.

8. Trokter M, Waksman G. Translocation through the conjugative type IV secretion system requires unfolding of its protein substrate. J Bacteriol. 2018;200: e00615–17.

9. Valenzuela-Gómez F, Arechaga I, Cabezón E. Nanopore sensing reveals a preferential pathway for the co-translocational unfolding of a conjugative relaxase–DNA complex. Nucleic Acids Res. 2023;51: 6857–6869.

10. Alonso-Lerma B, Jabalera Y, Samperio S, Morin M, Fernandez A, Hille LT, et al. Evolution of CRISPR-associated endonucleases as inferred from resurrected proteins. Nat Microbiol. 2023;8: 77–90.

11. Chen W, Ma J, Wu Z, Wang Z, Zhang H, Fu W, et al. Cas12n nucleases, early evolutionary intermediates of type V CRISPR, comprise a distinct family of miniature genome editors. Mol Cell. 2023;83: 2768–2780.e6.

12. Cui L, Bikard D. Consequences of Cas9 cleavage in the chromosome of *Escherichia coli*. Nucleic Acids Res. 2016;44: 4243–4251.

13. Trieu-Cuot P, Carlier C, Martin P, Courvalin P. Plasmid transfer by conjugation from *Escherichia coli* to Gram-positive bacteria. FEMS Microbiol Lett. 1987;48: 289–294.

14. Encinas D, Pilar Garcillán-Barcia M, Santos-Merino M, Delaye L, Moya A, De La Cruz F. Plasmid conjugation from proteobacteria as evidence for the origin of xenologous genes in cyanobacteria. J Bacteriol. 2014;196: 1551–1559.

15. Samperio S, Guzmán-Herrador DL, May-Cuz R, Cruz Martín M, Álvarez MA, Llosa M, et al. Conjugative DNA transfer from *E. coli* to transformation-resistant lactobacilli. Front Microbiol. 2021;12: 213.

16. Sadler JR, Tecklenburg M, Betz JL. Plasmids containing many tandem copies of a synthetic lactose operator. Gene. 1980;8: 279–300.

17. Killmann H, Benz R, Braun V. Properties of the FhuA channel in the Escherichia coli outer membrane after deletion of FhuA portions within and outside the predicted gating loop. J Bacteriol. 1996;178: 6913–6920.

18. Simon R, Priefer U, Pühler A. A broad host range mobilization system for in vivo genetic engineering: transposon mutagenesis in Gram negative bacteria. Nat Biotechnol. 1983;1: 784–791.

19. Gibson DG, Young L, Chuang RY, Venter JC, Hutchison 3rd CA, Smith HO. Enzymatic assembly of DNA molecules up to several hundred kilobases. Nat Methods. 2009;6: 343– 345.

20. Llosa M, Bolland S, de la Cruz F. Genetic organization of the conjugal DNA processing region of the IncW plasmid R388. J Mol Biol. 1994;235: 448–464.

21. Vieira J, Messing J. The pUC plasmids, an M13mp7-derived system for insertion mutagenesis and sequencing with synthetic universal primers. Gene. 1982;19: 259–268.

22. Lessl M, Balzer D, Weyrauch K, Lanka E. The mating pair formation system of plasmid RP4 defined by RSF1010 mobilization and donor-specific phage propagation. J Bacteriol. 1993;175: 6415–6425.

23. Grandoso G, Llosa M, Zabala JC, de la Cruz F. Purification and biochemical characterization of TrwC, the helicase involved in plasmid R388 conjugal DNA transfer. Eur J Biochem. 1994;226: 403–412.

24. Volke DC, Orsi E, Nikel PI. Emergent CRISPR–Cas-based technologies for engineering non-model bacteria. Curr Opin Microbiol. 2023;75: 102353.

25. Schmitz DJ, Ali Z, Wang C, Aljedaani F, Hooykaas PJJ, Mahfouz M, et al. CRISPR/Cas9 mutagenesis by translocation of Cas9 protein into plant cells via the *Agrobacterium* type IV secretion system. Front Genome Ed. 2020;2: 6.

